# Human pulmonary artery endothelial cells upregulate ACE2 expression in response to iron-regulatory elements: potential implications for SARS-CoV-2 infection of vascular endothelial cells

**DOI:** 10.1101/2021.04.08.437687

**Authors:** Quezia K Toe, Theo Issitt, Abdul S Mahomed, Ioannis Panselinas, Fatma Almaghlouth, Anne Burke-Gaffney, S John Wort, Gregory J Quinlan

## Abstract

Emerging studies from the ongoing covid-19 pandemic have implicated vascular dysfunction and endotheliitis in many patients presenting with severe disease. However, there is limited evidence for the expression of ACE2 (the principal co-receptor for Sars-Cov-2 cellular attachment) in vascular endothelial cells which has prompted the suggestion that the virus does not infect these cell types. However, the studies presented here demonstrate enhanced expression of ACE2 at the level of both mRNA and protein, in human pulmonary artery endothelial cells (PAECs) challenged with either IL-6 or hepcidin. Notably elevated levels both these iron-regulatory elements have been described in Covid-19 patients with severe disease and are further associated with morbidity and mortality. Additionally, levels of both IL-6 and hepcidin are often elevated in the elderly and in chronic disease settings, these populations being at greater risk of adverse outcomes from Sars-Cov-2 infection. A role for IL-6 and hepcidin as modulators of ACE2 expression seems plausible, additional, studies are required to determine if viral infection can be demonstrated in PAECs challenged with either of these iron-regulatory elements.

---

The current Covid-19 pandemic has resulted in significant global impacts for healthcare systems and beyond. Characterised as a respiratory infection, which for the majority of those infected is minor or asymptomatic, there are nevertheless significant life-threatening complications for a proportion of individuals. Age and or co-morbidities which are often chronic in nature are the main risk factors for severe Covid-19 infection with strong further associations with mortality. Severe acute respiratory distress syndrome (ARDS) is the primary disease presentation for those requiring intensive care treatment including the need for supportive ventilation or, in extreme cases, extracorporeal membrane oxygenation (ECMO). Whilst respiratory failure is a principal mortality factor for Covid-19, emerging data indicates that other organ failures including the kidney and cardiovascular system contribute to death. Evidence of widespread thrombosis with micro-thrombosis seen in the vasculature of lungs, kidneys and brain and other organs together with the presence of biomarkers of vascular dysfunction in blood samples from severe Covid-19 patients indicate a primary role for the endothelium in the disease process. It is recognised that SARS-CoV-2 primarily infect cells by using a viral surface glycoprotein (spike protein) which binds to angiotensin-converting enzyme related carboxypeptidase (ACE2) on the target cell to gain entry. ACE2 is reported to be expressed on many cell types including lung epithelia and vascular endothelium as reported in the previous SARS-CoV-1 epidemic^1^. However, current research efforts are unable to demonstrate any or very low levels of ACE2 expression in human endothelial cells^2^, which poses a question as to how SARS-CoV-2 can infect and cause endotheliitis as widely reported. IL-6 is reported to be greatly elevated in patients with severe Covid-19 infection^3^. As a key positive regulator for hepcidin biosynthesis this may suggest impacts for iron metabolism in these patients; elevated serum ferritin further reinforces this notion. Hepcidin, described as the global regulator for iron homeostasis, is released from the liver and binds to the cellular iron exporter, ferroportin, causing internalisation; thereby, preventing cellular iron export. The axis is operational in the liver, gut and cells chiefly involved in iron recycling for biosynthetic processes including erythropoiesis (reviewed by Reichert *et al* 2017)^4^. Our recent studies have identified a localised hepcidin/ ferroportin axis that is operational in human pulmonary artery smooth muscle cells (hPASMCs); importantly, IL-6 promotes hepcidin production by hPASMCs^5^. We now report the presence of this axis is human pulmonary artery endothelial cells (hPAECs) (Fig. 1A), which shows ferroportin expression and down-regulation by hepcidin (confocal images) together with a western blot which further demonstrates IL-6 mediated down-regulation. These observations prompted us to consider a role for IL-6 and the hepcidin/ferroportin (H/F) axis as a modular for ACE2 expression in human pulmonary artery endothelial cells (hPAECs), given that interactions between ACE2 and IL-6 have previously been described^6^. Studies were undertaken with hPAECs obtained from ATCC^®^.

**Figure.**
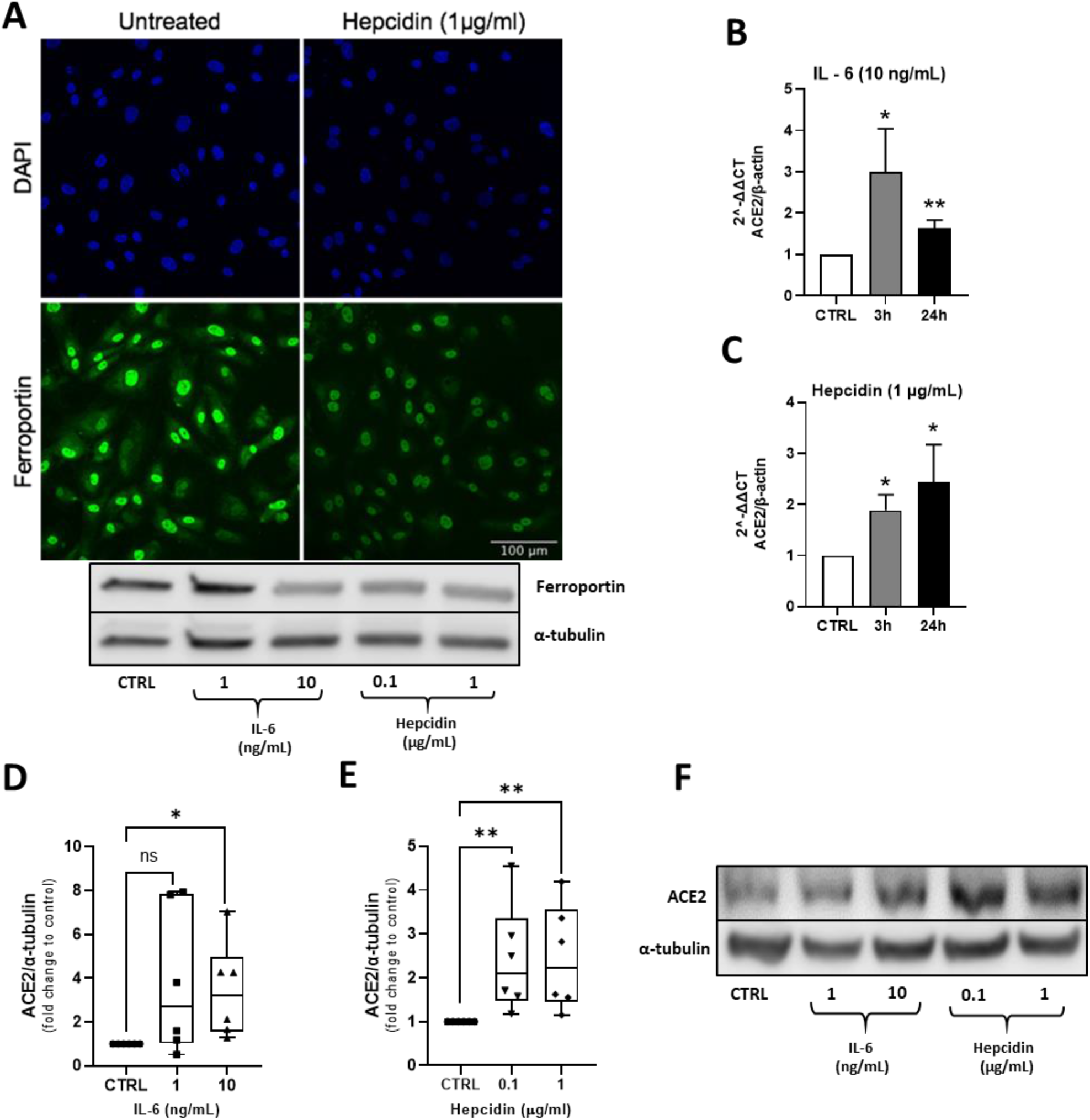
Hepcidin and IL-6 upregulate ACE2 expression in hPAECs. **(A)** *Ferroportin is expressed by hPAECs and modulated by IL-6 and hepcidin*. hPAECs were either untreated (CTRL) or incubated with 1 μg/mL hepcidin for 24 h. Cells were then fixed using 4% paraformaldehyde, permeablised with 0.2% triton x-100, blocked in 10% BSA and incubated with rabbit anti-ferroportin (1:1000), washed with PBS 3 times, incubated with Alexa Fluor anti-rabbit 700 (1:500), counterstained with DAPI and visualised. Images show maximum intensity per pixel across the Z plane. hPAECs were treated with media alone, IL-6 (1 and 10 ng/mL) and hepcidin (0.1 and 1 μg/mL) for 24 h, cells were lysed, and 50 ug of protein separated on 4-15% SDS-page and transferred onto nitrocellulose membranes. Western blotting was performed using rabbit anti-Ferroportin (1:1000; NBP1-21502, NOVUS) and α-tubulin (1:2000; 3873S, Cell Signalling). **(B-C)** *ACE2 mRNA transcription was upregulated by IL-6 and hepcidin*. RT-PCR was performed using SensiFast SYBR green Lo-ROX (Meridian Biosciences) with human ACE2 primers (forward – GTTTGTAACCCAGATAATCCAC; reverse – AATGATTTGCTCTTGCCATC) and β-actin (forward – GACGACATGGAGAAAATCTG; reverse – ATGATCTGGGTCATCTTCTC), after hPAECs were treated with media alone, IL-6 (10 ng/mL) or hepcidin (μg/mL) for 3 or 24h. The values were normalised as fold changes to the control of untreated cells at each corresponding time point. N=3, student’s t test was performed; *p<0.05, **p<0.01. **(D, E, F)** *IL-6 and hepcidin increased ACE2 protein expression*. hPAECs were treated with media alone, IL-6 (1 and 10 ng/mL) and hepcidin (0.1 and 1 μg/mL) for 24 h, cells were lysed, and 50 ug of protein separated on 4-15% SDS-page and transferred onto nitrocellulose membranes. Western blotting was performed using rabbit anti-ACE2 (1:500; ab15348, Abcam) and α-tubulin (1:2000; 3873S, Cell Signalling). Relative protein expression quantification was performed with ImageJ software. Values were normalised to α-tubulin and fold change calculated against control. Three hPAEC donors were used, two experiments were run for each donor. One-way ANOVA non-parametric Kruskal-Wallis tests were performed with post-hoc Dunn’s test for multiple testing correction; *p<0.05, **p<0.01.

When challenged with either IL-6 or hepcidin, significant upregulation of ACE2 mRNA over time was observed with hPAECs from 3 donors (Fig. 1B and 1C). Importantly, we also observed elevated levels of ACE2 protein expression by western blot over time (Fig. 1C and 1D). These observations may provide an explanation for viral vascular endothelial cell infection seemly seen in severe Covid-19 disease. Given the importance of shear stress as a modulator of gene expression for hPAECs in vivo, an assessment of ACE2 transcript expression was undertaken by subjecting hPAECs to laminar flow (15 dyn/cm^2^) in a parallel-plate fluid flow chamber. However, ACE2 did not prove to be a flow responsive gene (data not shown).

It is noteworthy that key aspects of the ageing process, inflammaging^7^ and the anaemia of ageing^8^, are associated with elevated levels of IL-6 and hepcidin. Similar effects are also seen in chronic/comorbid disease settings (anaemia of chronic disease)^9^. Given our findings, it is not unreasonable to suggest that ageing and or chronic disease may predispose these groups to more severe vascular disease because of the likelihood of elevated ACE2 expression linked to elevated IL-6/hepcidin. Emerging literature is also establishing links between the hepcidin/ferroportin axis and SARS-CoV-2^10^ with some sequence homology between the SARS-CoV-2 spike protein and hepcidin reported^11^. Interestingly, CD26 another suggested co-receptor for SARS-CoV-2 is also known to bind to ferroportin^12^. Moreover, published studies from China and Italy^13,14^ have reported strong associations with severity of disease and mortality and hepcidin levels. The significance of these findings remains unclear but does further highlight a potential link between dysregulated iron metabolism and Covid-19 infection.

Whilst our studies demonstrate upregulation of ACE2 in human hPAECs in response to iron regulatory elements, studies need to be undertaken to establish if such effects result in enhanced SARS-CoV-2 infection. Should this be the case, targeting the H/F axis may well provide therapeutic benefit in severe Covid-19 infection and further supports the use of the IL-6 receptor antibody, tocilizumab^15^.

Authors wish to thank the British Heart Foundation for grant support.

## References

1. Hamming, I., Timens, W., Bulthuis, M., Lely, A., Navis, G. and van Goor, H., 2004. Tissue distribution of ACE2 protein, the functional receptor for SARS coronavirus. A first step in understanding SARS pathogenesis. The Journal of Pathology, 203(2), 631–637.

2. McCracken, I., Saginc, G., He, L., Huseynov, A., Daniels, A., Fletcher, S., Peghaire, C., Kalna, V., Andaloussi-Mäe, M., Muhl, L., Craig, N., Griffiths, S., Haas, J., Tait-Burkard, C., Lendahl, U., Birdsey, G., Betsholtz, C., Noseda, M., Baker, A. and Randi, A., 2021. Lack of Evidence of ACE2 Expression and Replicative Infection by SARSCoV-2 in Human Endothelial Cells. Circulation, 143(8), 865–868.

3. Han, H., Ma, Q., Li, C., Liu, R., Zhao, L., Wang, W., Zhang, P., Liu, X., Gao, G., Liu, F., Jiang, Y., Cheng, X., Zhu, C. and Xia, Y., 2020. Profiling serum cytokines in COVID-19 patients reveals IL-6 and IL-10 are disease severity predictors. Emerging Microbes & Infections, 9(1), pp.1123–1130.

4. Reichert, C., da Cunha, J., Levy, D., Maselli, L., Bydlowski, S. and Spada, C., 2017. Hepcidin: Homeostasis and Diseases Related to Iron Metabolism. Acta Haematologica, 137(4), 220–236.

5. Ramakrishnan, L., Pedersen, S., Toe, Q., West, L., Mumby, S., Casbolt, H., Issitt, T., Garfield, B., Lawrie, A., Wort, S. and Quinlan, G., 2018. The Hepcidin/Ferroportin axis modulates proliferation of pulmonary artery smooth muscle cells. Scientific Reports, 8(1), 12972.

6. Mokuda, S., Tokunaga, T., Masumoto, J. and Sugiyama, E., 2020. Angiotensin-converting Enzyme 2, a SARS-CoV-2 Receptor, Is Upregulated by Interleukin 6 through STAT3 Signaling in Synovial Tissues. The Journal of Rheumatology, 47(10), 1593–1595.

7. Franceschi, C. and Campisi, J., 2014. Chronic Inflammation (Inflammaging) and Its Potential Contribution to Age-Associated Diseases. The Journals of Gerontology Series A: Biological Sciences and Medical Sciences, 69(S1), S4–S9.

8. Halawi, R., Moukhadder, H. and Taher, A., 2017. Anemia in the elderly: a consequence of aging?. Expert Review of Hematology, 10(4), 327–335.

9. Madu, A. and Ughasoro, M., 2016. Anaemia of Chronic Disease: An In-Depth Review. Medical Principles and Practice, 26(1), 1–9.

10. Cavezzi, A., Troiani, E. and Corrao, S., 2020. COVID-19: Hemoglobin, Iron, and Hypoxia beyond Inflammation. A Narrative Review. Clinics and Practice, 10(2), 24–30.

11. Ehsani, S., 2020. COVID-19 and iron dysregulation: distant sequence similarity between hepcidin and the novel coronavirus spike glycoprotein. Biology Direct, 15(1).

12. Raha, A., Chakraborty, S., Henderson, J., Mukaetova-Ladinska, E., Zaman, S., Trowsdale, J. and Raha-Chowdhury, R., 2020. Investigation of CD26, a potential SARS-CoV-2 receptor, as a biomarker of age and pathology. Bioscience Reports, 40(12), BSR20203092.

13. Zhou, C., Chen, Y., Ji, Y., He, X. and Xue, D., 2020. Increased Serum Levels of Hepcidin and Ferritin Are Associated with Severity of COVID-19. Medical Science Monitor, 26.

14. Nai, A., Lorè, N., Pagani, A., De Lorenzo, R., Di Modica, S., Saliu, F., Cirillo, D., Rovere-Querini, P., Manfredi, A. and Silvestri, L., 2020. Hepcidin levels predict Covid- 19 severity and mortality in a cohort of hospitalized Italian patients. American Journal of Hematology, 96(1).

15. Xu, X., Han, M., Li, T., Sun, W., Wang, D., Fu, B., Zhou, Y., Zheng, X., Yang, Y., Li, X., Zhang, X., Pan, A. and Wei, H., 2020. Effective treatment of severe COVID-19 patients with tocilizumab. Proceedings of the National Academy of Sciences, 117(20), 10970–10975.

